# Cell type specific CaMKII activation patterns revealed by CaMKAR, a bioactivity reporter deployable in living cells

**DOI:** 10.1101/2025.06.11.658946

**Authors:** Alex Severino, Oscar E Reyes Gaido, Bian Liu, Gabriel Lopez-Cecetaite, An-Chi Wei, Giovanni Rosales-Soto, Erick O Hernandez-Ochoa, Elizabeth D Luczak

## Abstract

An accurate and precise mechanism for measuring CaMKII activity in living cells is invaluable in the search for effective and targeted CaMKII-based therapeutics. Here, we employ our recently published CaMKII Activity Reporter (CaMKAR) biosensor in order to investigate the spatiotemporal dynamics of CaMKII activation in three different types of cells – cardiac myocytes, skeletal myocytes, and neurons. In doing so, we found a greater rate of CaMKII activation in skeletal muscle compared to cardiac muscle and also delineated CaMKAR’s ability to measure discrete CaMKII activation events in the presence of individual action potentials. By modifying the original CaMKAR sequence, we generated sensors that can be localized to subcellular compartments and thereby preferentially detect the activity of specific spatially-distributed CaMKII isoforms. Finally, we utilized the live-cell data to generate mathematical models of CaMKII activation kinetics, both as an integrated function across multiple calcium transients and as discrete on-off events following individual depolarizations. By furthering our understanding of CaMKII activity profiles across cell types and within subcellular compartments, we hope to support development of CaMKII inhibitors that are optimally precise and potent.

## Introduction

The multifunctional Ca^2+^ and calmodulin-dependent protein kinase II (CaMKII) dynamically couples intracellular Ca^2+^ activity to fundamental cellular responses. Across various types of excitable cells, this signal transduction is crucial for physiologic processes. In neurons, long-term potentiation requires CaMKII recruitment to the postsynaptic density, where CaMKII aids in restricting the movement of AMPA receptors in order to amplify incoming stimuli [1–3]. In cardiac muscle, CaMKII is an important regulator of pacemaker physiology in the sinoatrial node [4, 5] and energy metabolism [6]. In skeletal muscle, CaMKII plays a role in maintaining tension during muscle contraction as well as contraction-induced glucose uptake [7, 8].

In mammals, CaMKII is expressed from 4 genes (α, β, γ, δ) that are alternatively spliced, leading to tissue specificity and subcellular localization. CaMKII is a heteromultimer with each monomer containing an N-terminal catalytic domain, a regulatory domain, and a C-terminal association domain [4] (4). Canonical activation of CaMKII occurs in excitable cells when cells depolarize and undergo intracellular calcium influx. The calcium then binds to calmodulin (CaM), and calcified CaM binds to and activates CaMKII. Sustained intracellular calcium leads to autophosphorylation of the enzyme at the T287 residue [9]. T287 autophosphorylation allows CaMKII to become autonomously active, independent of upstream calcium signaling [10]. Additional activating post-translational modifications such as oxidation, nitrosylation, and O-GlcNAcylation enable CaMKII to undergo mechanistic tuning of its activation profile depending on the local cellular environment.

While CaMKII signaling regulates various physiologic processes, hyperactivation of CaMKII contributes to cellular dysfunction and is a known driver of multiple pathologies. The phenomenon of CaMKII-induced dysfunction is perhaps best characterized in the cardiovascular system, where CaMKII hyperactivation leads to endothelial dysfunction, structural heart disease, and arrythmias [4]. CaMKII inhibitors have successfully been developed, however, they have yet to translate to clinical benefits.

One outstanding limitation to CaMKII inhibition is the detriment of potential off-target effects. Given CaMKII’s important role in neuronal long-term potentiation, indiscriminate inhibition would conceivably risk impairment of learning and memory [4]. Further investigation of isoform-and tissue-specific CaMKII characteristics will therefore be critical to the development of clinically viable enzyme inhibitors.

Despite the importance of CaMKII in health and disease and our need for a precise understanding of its activation patterns, kinetic measurements of CaMKII activity were previously lacking in living cells. In order to advance our ability to measure CaMKII precisely and efficiently, we recently developed a new CaMKII Activity Reporter (CaMKAR). CaMKAR is a real-time, reversible, fluorescence-based biosensor of CaMKII enzymatic activity, reporting direct substrate phosphorylation with a superior signal:noise and temporal response compared to previous biosensors. Using cp-GFP, CaMKAR possesses the ability to generate both ratiometric and intensiometric readout of CaMKII activation in living cells [11].

Here, we employ CaMKAR to quantifiably demonstrate CaMKII activation kinetics across different biological systems. We measured CaMKAR signal in real time in three types of excitable cells: cardiomyocytes, skeletal myofibers, and neurons. Our results demonstrate, for the first time, that CaMKII operates over a broad range of temporal dynamics in different cell types, providing valuable insight to our understanding of this fundamental molecular signal important for human pathophysiology. We highlight CaMKAR’s ability to achieve targeted subcellular localization, validating our sensor as a tool to further elucidate CaMKII’s spatiotemporal profile in future studies. Finally, we utilize nonlinear ordinary differential equations (ODEs) in order to mathematically model the activation of CaMKII in these different systems.

## Results

In order to assess CaMKAR’s kinetic ability to precisely delineate temporal differences in CaMKII activation, we coexpressed CaMKAR and the genetically encoded calcium indicator jRCaMP1a in HEK293T cells. This system allows us to simultaneously quantify and track both CaMKAR phosphorylation and calcium influx. We treated these cells with ionomycin, a calcium ionophore, and observed a rise in CaMKAR fluorescence beginning only 4 seconds after initial calcium entry (Fig 1). With this, we verify that CaMKAR is sensitive to fluctuation in CaMKII activity on the timescale of seconds, supporting the sensor’s ability to detect meaningful distinctions that may exist between different cell populations.

**Figure 1.**
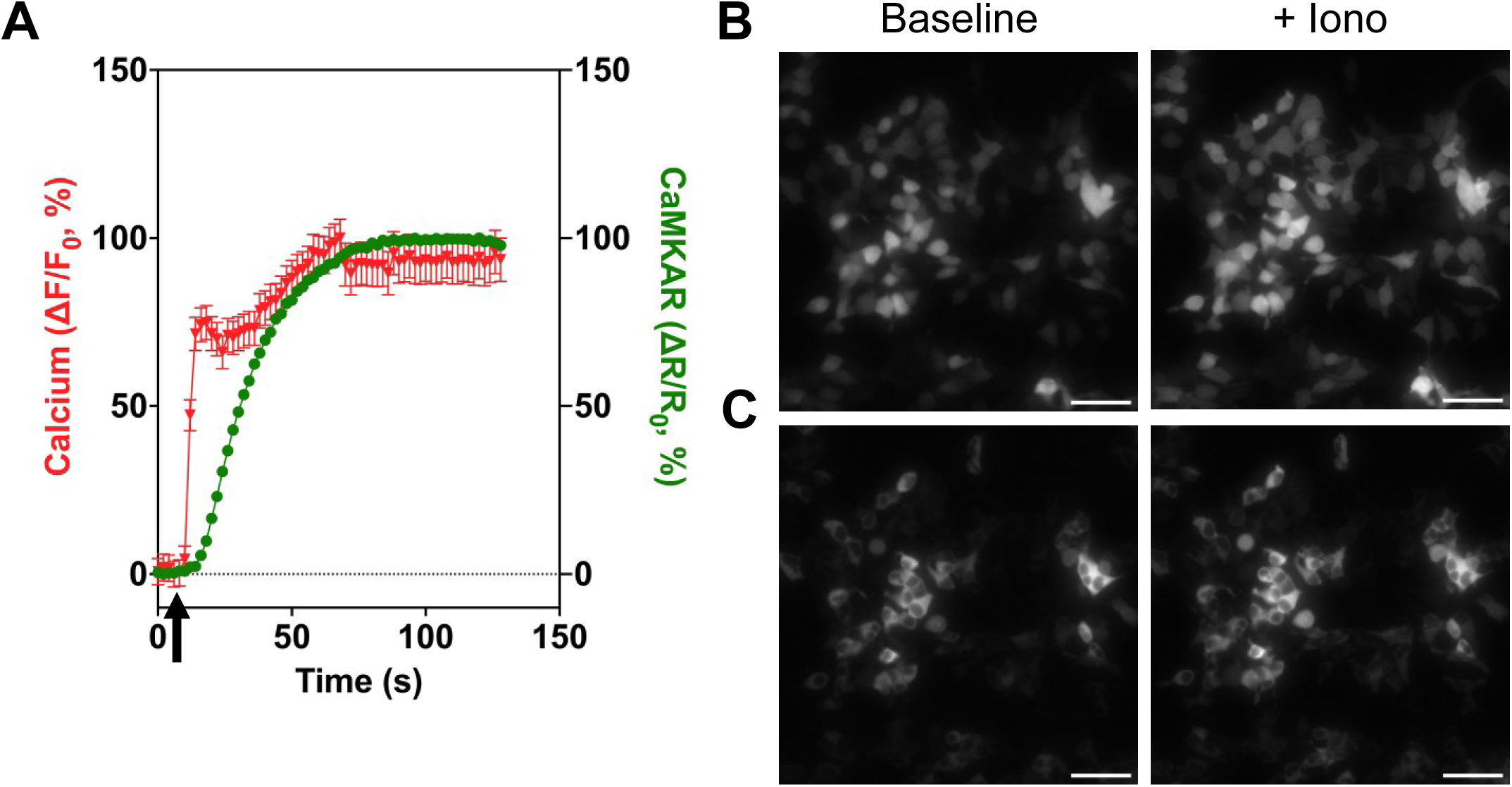
Ca^2+^ and CaMKAR fluorescence in HEK293T cells treated with ionomycin. (**A**) Fluorescence signal of Ca^2+^ (jRCaMP1a) and CaMKAR measured simultaneously in HEK293T cells after administration of calcium ionophore, ionomycin (iono), at time t=10 sec (arrow). Both signals normalized to 100% of maximum intensity with data plotted as mean ± SEM. Representative fluorescence microscopy images of HEK293T cells transfected with jRCaMP1a (**B**) and CaMKAR (**C**) plasmid before (left) and after (right) addition of ionomycin. Scale bars, 50µm.

We next utilized a sensor-encoding adenovirus to express CaMKAR in neonatal rat ventricular myocytes (NRVMs). We measured CaMKAR signal at baseline and in the presence of two validated activators of CaMKII (Fig 2A), electrical pacing and angiotensin II (AngII) (Fig 2B). Electrical field pacing of cardiac myocytes induces calcium transients that stimulate CaMKII activity [12, 13]. The amplitude of change in CaMKAR fluorescence was nearly 4x greater in cells stimulated at 2Hz versus 1 Hz while the time to half peak remained similar (Table 1). These data illustrate the dependence of CaMKII activation amplitude on the frequency of stimulation [14]. AngII, on the other hand, has been shown to activate CaMKII through oxidation of CaMKII secondary to increased ROS production [15–17]. AngII stimulation produced peak amplitude similar to 1Hz electrical stimulation, but had a faster upward slope and time to half peak. (Table 1). Taken together, the data show the versatility of the CaMKAR sensor in measuring CaMKII activation via alternative pathways.

**Fig. 2.**
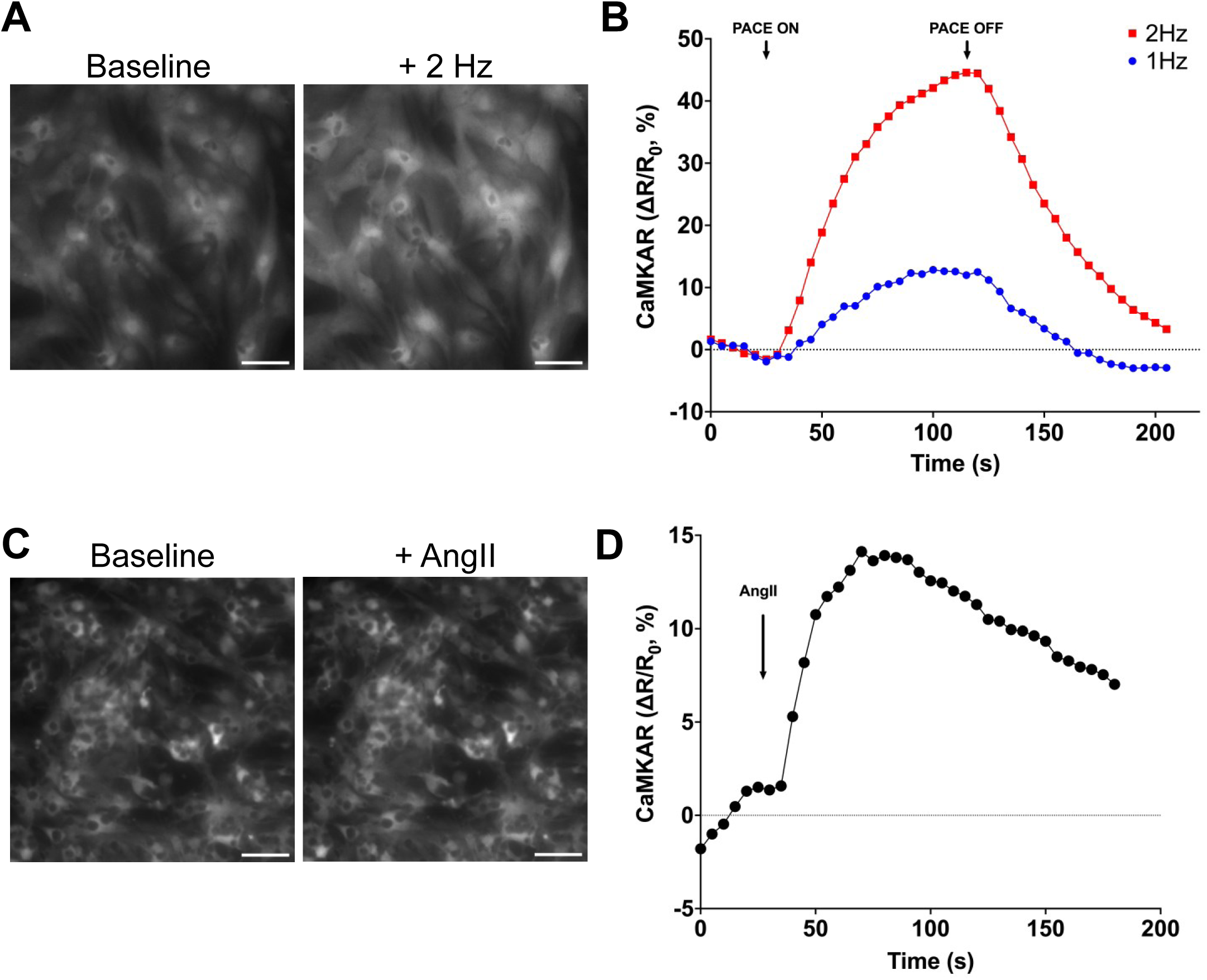
CaMKAR activation in NRVMs. (**A**) Representative fluorescence microscopy images of neonatal rat ventricular myocytes (NRVMs) infected with CaMKAR adenovirus before (left) and after (right) electrical stimulation at 2 Hz. (**B**) Quantification of ratiometric CaMKAR fluorescence in NRVMs sampled every 5 seconds is shown during 1 and 2 Hz pacing, with change in fluorescence at each timepoint (ΔR) normalized to the baseline (R_0_) to produce ΔR/R_0_. (**C**) Representative fluorescence microscopy images of NRVMs infected with CaMKAR adenovirus before (left) and after (right) treatment with AngII. (**D**) Quantification of ratiometric CaMKAR fluorescence in NRVMs following AngII treatment (arrow). Ratiometric signals are normalized to baseline fluorescence. Scale bars, 50µm.

**Table 1.** Kinetics of CaMKII across cell types. Calculations performed using timelapse curves in Figures 2, 3, and 4. Time to half-peak defined as time elapsed in seconds from beginning of stimulation to 50% of the peak fluorescence signal. Peak amplitude is reported as the fold-change between the maximum fluorescence in each cell type compared to baseline. Upward slope calculated in the linear portion of each curve.

| | | Time to half-peak (s) | Upward slope (($\Delta R/R_0$)/s) | Peak amplitude ($\Delta R/R_0$ ,%) |
| --- | --- | --- | --- | --- |
| Cardiomyocytes | 1Hz | 31.75 | 0.32 ± 0.019 | 12.84 |
|  | 2Hz | 28.53 | 0.99 ± 0.031 | 44.58 |
|  | AngII | 17.67 | 0.61 ± 0.037 | 14.13 |
| Skeletal Muscle | 10 Hz | 10.03 | 1.41 ± 0.015 | 24.16 |
|  | 70 Hz | 7.34 | 3.81 ± 0.046 | 38.96 |
|  | 140 Hz | 11.77 | 2.11 ± 0.029 | 40.00 |
| Neurons | Glycine | 0.43 | 31.46 ± 5.58 | 13.63 |

In skeletal muscle, CaMKII is known to regulate excitation-transcription coupling and mediate the muscular response to exercise [7, 18]. In skeletal muscle fibers isolated from the flexor digitorum brevis (FDB) of mice, we introduced CaMKAR using electroporation and activated CaMKII with electrical pacing. We electrically stimulated these fibers at 10, 70, and 140 Hz to fully capture the typical range of frequencies for both slow-twitch and fast-twitch fibers and measured CaMKAR intensiometrically to allow for faster acquisition of signal readout (Fig 3A-B). The amplitude of change in fluorescence was similar and greatest during 70 Hz and 140 Hz pacing compared 10 Hz (Table 1). The rate of activation (time to half peak and upward slope), however, was fastest in the 70Hz condition.

**Figure 3.**
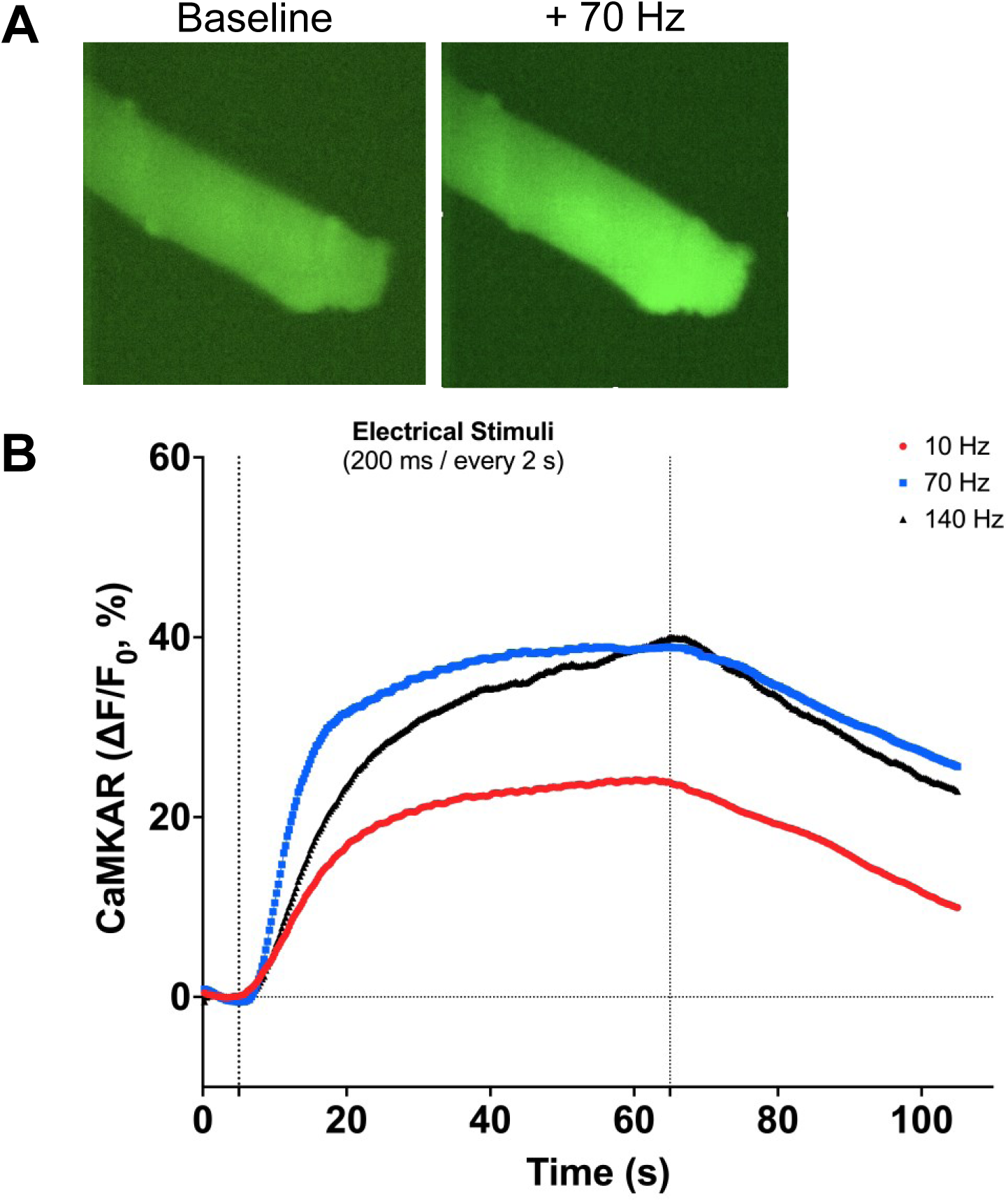
CaMKAR activation in skeletal muscle. (**A**) Representative fluorescence images of CaMKAR in mouse flexor digitorum brevis (FDB) skeletal muscle fibers at 488 nm before (left) and after (right) electrical stimulation for 60 seconds at 70 Hz (500 ms train / every 2 sec). (**B**) Quantification of change in fluorescence (ΔF) normalized to baseline signal (F_0_) with pacing at 10 Hz, 70 Hz, and 140 Hz. N = 4 animals, 7-13 fibers considered for each condition in separate experiments.

Up until this point the kinetics of CaMKII have perhaps most extensively been modeled in neurons due to CaMKII’s function as a bistable switch in long-term potentiation and memory formation [19, 20]. It has therefore been demonstrated using previous FRET-based CaMKII sensors that CaMKII signaling occurs rapidly within dendritic spines of neurons, on the order of milliseconds [21]. Here, we expressed CaMKAR in rat hippocampal neurons and stimulated with glycine in order to assess our sensor’s ability to detect CaMKII activation on a rapid timescale. We simultaneously imaged calcium transients during neuronal excitation using the jRGECO1a calcium sensor in order to directly correlate CaMKAR signal to individual calcium transients (Fig 4A-B). In this way, we utilize CaMKAR to measure discrete CaMKII activation events, compared to the integration of multiple action potentials over a longer duration of time that are measured in our paced cardiomyocytes. In these CaMKAR neurons, fluorescent signal reached half-peak intensity in 0.43 seconds with an upward slope of 31.46 ± 5.58 units per second (Table 1). Furthermore, the time to half-peak of the CaMKAR signal trailed that of the dendritic calcium transient by 100 milliseconds (Fig 4C), emphasizing CaMKII’s rapid sensitivity to alterations in intracellular calcium.

**Figure 4.**
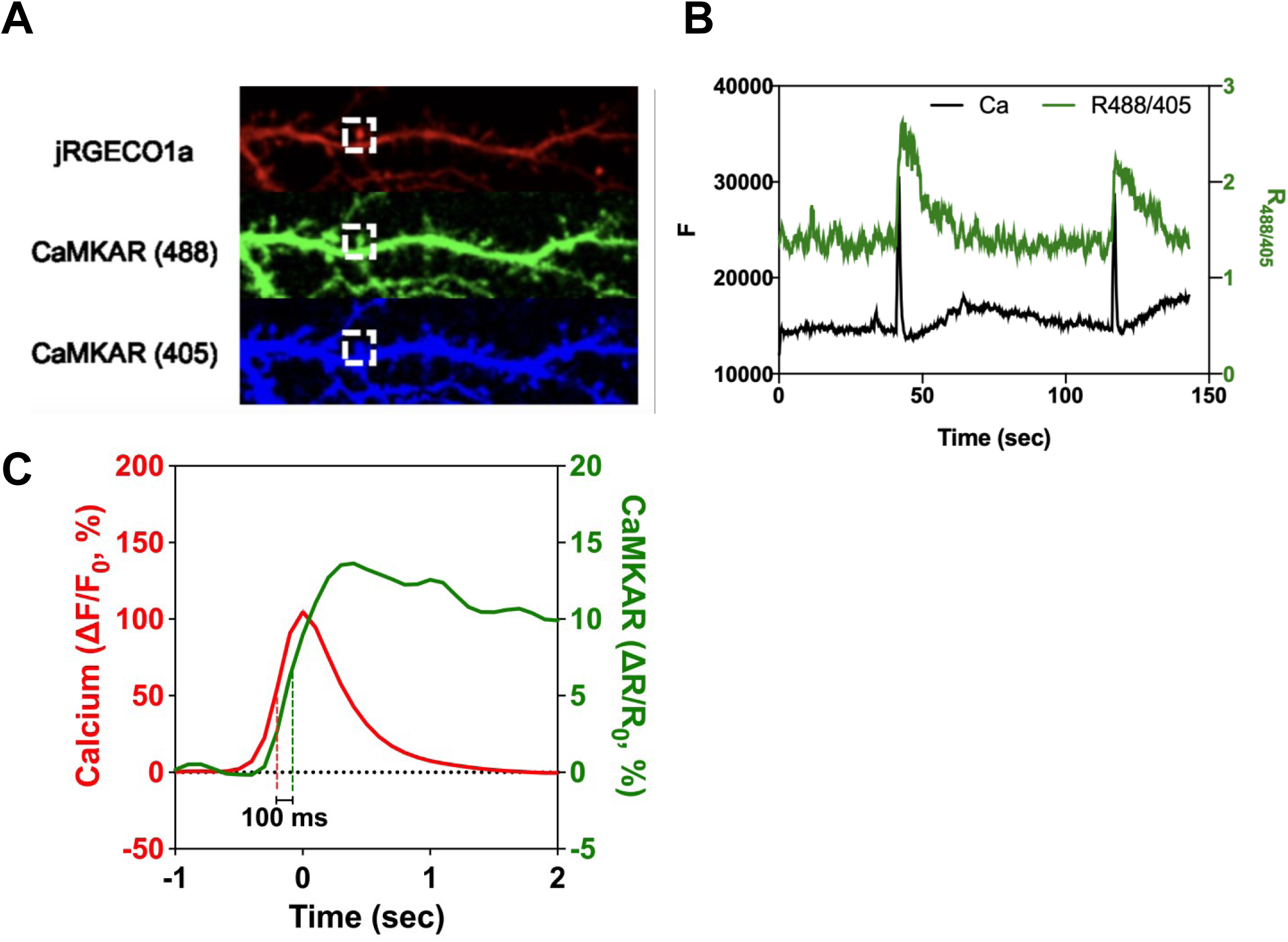
**CaMKAR activation in neurons**. Primary mouse neurons were made to express CaMKAR and the jRGECO1a Ca^2+^ via transfection. Single pulses of field stimulation induced CaMKII activity. (**A**) Representative fluorescence confocal images of dendritic spines of hippocampal neurons split by channel for excitation of CaMKAR at either 488 nm or 405 nm and jRGECO1a with excitation at 561 nm. (**B**) Time lapse summary data plotted as fluorescence units measured from single Z slice images (F) imaged at 10 Hz. White boxes on (A) denote the portion of image used for tracing in (B). (**C**) For a single transient event, simultaneous Ca^2+^ (red) and CaMKAR (green) tracings are shown with fluorescence normalized to baseline. Vertical lines denote half-peak for each tracing.

Beyond CaMKAR’s ability to effectively delineate the cell-specific timescales of CaMKII activity in living cells, it is also uniquely positioned as a genetically encoded sensor to be targeted to specific subcellular compartments. CaMKII has been shown to possess different characteristics and enact various effects depending on its localization. For instance, it has been shown that mitochondrial CaMKII may be necessary for pathologic development of dilated cardiomyopathy after myocardial infarction, while the nuclear variant CaMKIIδB appears to provide protection from the detrimental effects of cardiac ischemia/reperfusion [6, 22]. Therefore, it would greatly expand the utility of any CaMKII sensor to be able to report activity of subcellularly localized CaMKII.

In order to test CaMKAR in these settings, we appended different localization tags to the protein’s C terminus and expressed these targeted CaMKAR variants in HEK cells (Fig 5A). After verifying that CaMKAR could be successfully localized to the nucleus, the cytosol, and the mitochondria, we next sought to ensure that the sensor retained its reporting ability in each of these different environments. To do so, we took advantage of the localization of different CaMKII splice isoforms by introducing constitutively active CaMKII variants alongside compartmentalized CaMKAR. These variants included the cytosolic CaMKII δ9 and δC, the nuclear CaMKII δB, and an artificial variant that is trafficked to mitochondria through a mitochondrial targeting signal (MTS) fused to CaMKIIδ. Cytosolic CaMKAR reported a signal with each of the CaMKII variants (Fig 5B), consistent with current literature suggesting that each of these expressed variants can be enriched in the cytoplasm [22]. CaMKAR targeted to the nucleus exhibited the greatest fluorescence with the δB variant, while the MTS-δ fusion CaMKII protein generated the greatest signal above noise with mitoCaMKAR compared to other variants in mitochondria (Fig 5B-D).

**Figure 5.**
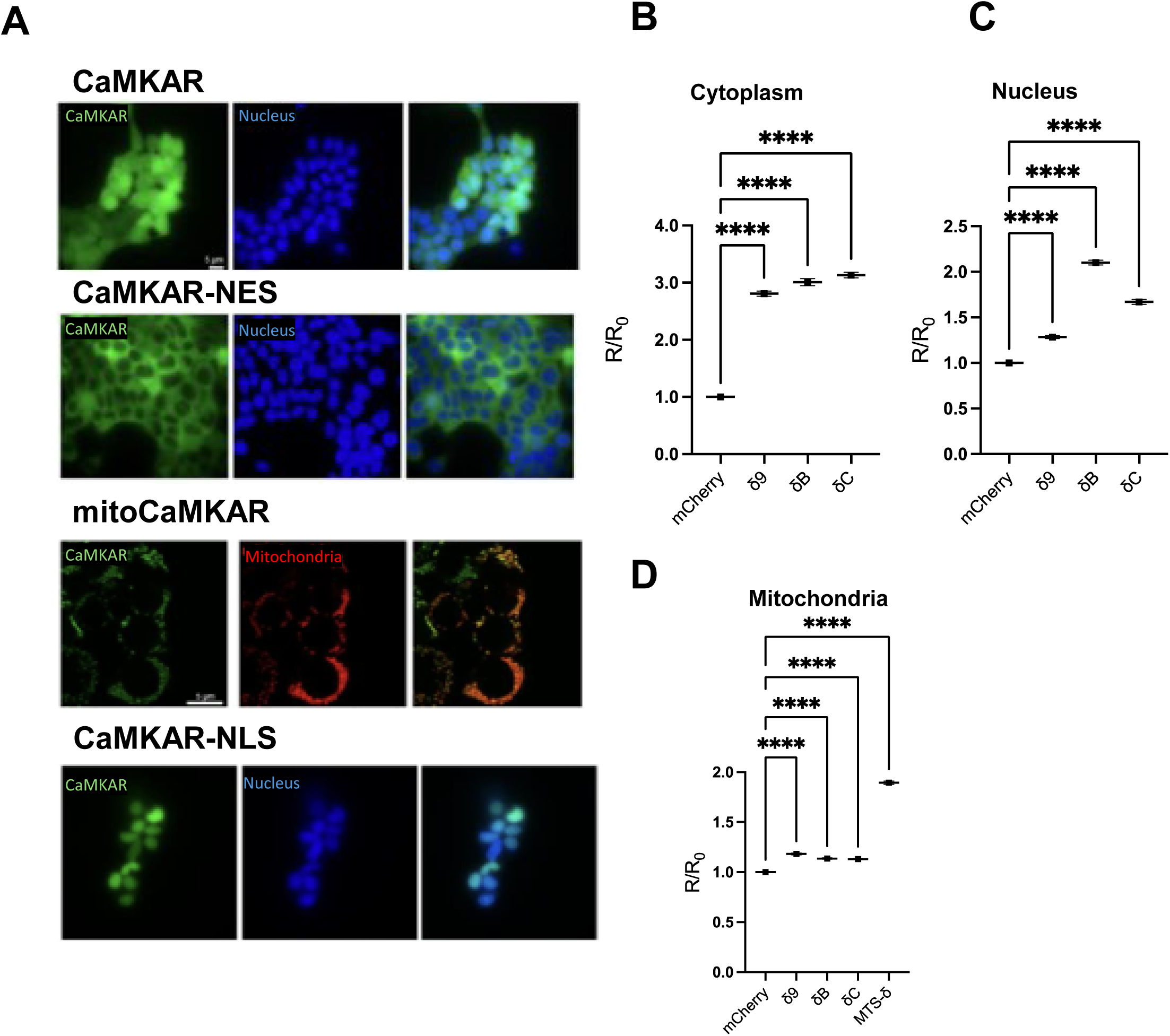
Subcellular compartmentalization of CaMKAR. Representative images are shown of CaMKAR localization modified by inclusion of nuclear export signal (NES), nuclear localization signal (NLS), and mitochondrial localization via mitochondrial targeting sequence (MTS, mito) (**A**). Nuclei were stained by Hoechst 33342 and mitochondria were stained with mitoTracker Far Red. CaMKAR-NES (**B**), CaMKAR-NLS (**C**), and mitoCaMKAR (**D**) were expressed in HEK293T cells with constitutively active CaMKII variants known to primarily reside in either cytosol (δ9 and δC), nucleus (δB), and mitochondria (MTS-δ fusion). Confocal imaging was used to derive CaMKAR ratio (R/R_0_). Data is shown as mean and error bars denote S.E.M from many cells across 3 biological replicates. Fluorescence across groups was compared using ordinary one-way ANOVA (**** = *p* < 0.0001)

Using temporal data generated from CaMKAR in 3 distinct cell types, we mathematically modeled CaMKII activation kinetics in each of these systems. The kinetic scheme represents the state transitions between calcium binding to calmodulin and binding to CaMKII (Fig. 6A). The model parameters were taken from the CaMKIIα model developed for neurons by Chang et al. [21]. The parameter set was tested in the neuron (Fig. 6B) under stimulations every 60sec. 1Hz and 2Hz of pacing were tested in NRVM and intermittent pulses with 10Hz, 70Hz, and 140Hz of the parameters were tested in skeletal muscle (Fig. 6B). Parameters of calmodulin affinity and autophosphorylation rate were adjusted based on the CaMKII isoform studies reported by Gaertner et al. [23, 24] and the experimental results in Figures 2, 3, and 4. CaMKIIα is the primary isoform in the brain, CaMKIIδ in the heart, and CaMKIIγ in the smooth muscle [25]. The calcium waveforms and CaM-CaMKII parameters (Supplemental Table 1) were adjusted to simulate the CaMKII activities measured by CaMKAR in these three cell types.

**Figure 6.**
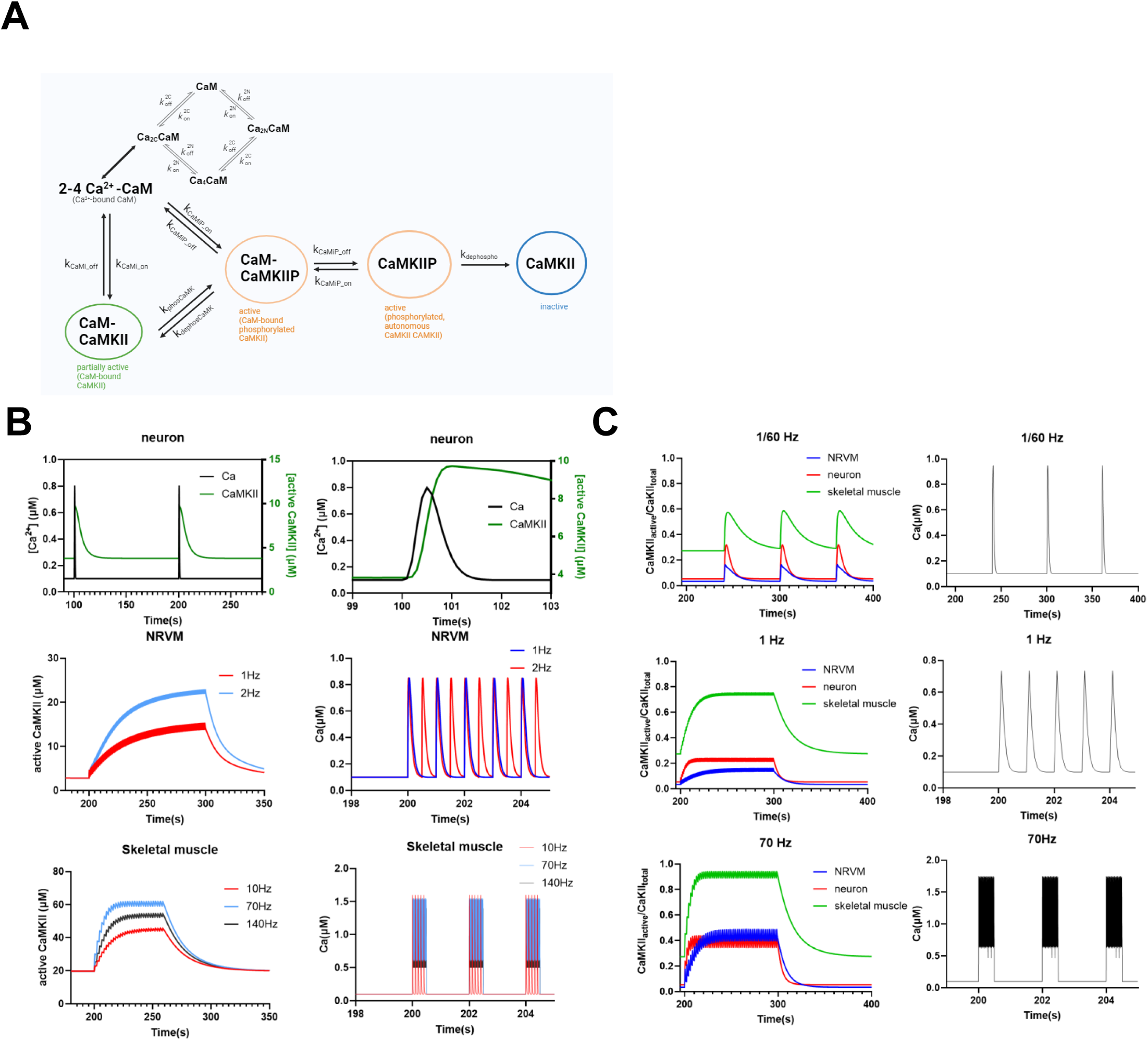
Simulations of CaMKII activities in different cell types using the kinetic model of Ca^2+^-CaM-CaMKII. (**A**) The schematic of interaction among Ca^2+^, calmodulin (CaM), and CaMKII. Calmodulin binds with 2–4 Ca^2+^ ions to form Ca-CaM complexes. CaMKII-P is the phosphorylated CaMKII. CaMKII denotes the inactive form after dephosphorylation. (**B**) Kinetic parameters and binding constants of CaMKII models were determined by fitting to the CaMKII activities measured using CaMKAR in neurons, skeletal muscle, and NRVM in Figures 2, 3, and 4. (**C**) Simulations of CaMKII activities under different pacing protocols (continuous pulses at 1/60 Hz, and 1Hz, and intermittent pulses at 70 Hz) for different cell types using the kinetics parameters obtained from (B). The active CaMKII was the sum of all the active CaMKII (CaMKIIP, CaM_CaMKIIP). The total CaMKII was the sum of all forms of CaMKII.

After finding the parameter sets for each cell type, we tested these three cell types’ CaMKII response under different pacing protocols as were used in the experiment (Fig. 6C). CaMKII activity rises during the calcium spike interval as calcium binds to CaM and CaMKII, and phosphorylation occurs. CaMKII activity then falls after the calcium pulse as CaM slowly dissociates and dephosphorylation occurs. At low pacing frequency, such as 1/60 Hz, CaMKII can completely go back to inactive form in each pulse. At the higher frequency of pacing (1Hz-140Hz), Ca^2+^-CaM bound molecules accumulate before dissociation, increasing CaM binding to neighboring subunits, allowing for autophosphorylation, and active forms of CaMKII to accumulate. After the calcium pulse stimulation ends, CaMKII gradually dephosphorylated, and the inactive unbound form of CaMKII accumulates.

Neurons show the fastest activation and accumulation of active CaMKII after stimulation. Skeletal muscle CaMKII shows slower kinetics in deactivation after stopping pacing.

## Discussion

In this study we deployed novel applications for our recently developed live cell CaMKII biosensor CaMKAR. We first demonstrated CaMKAR’s rapid detection of CaMKII activation using simultaneous measurement of CaMKAR and calcium signal in order to verify the tight temporal connection between calcium transients and CaMKAR fluorescence. We subsequently used CaMKAR to probe the spatiotemporal profile of CaMKII in excitable cells including cardiomyocytes, skeletal muscle fibers, and neurons. We found unique profiles of CaMKII activation patterns across these different cell types and with different mechanisms of stimulation within a cell type. For instance, the peak amplitude of CaMKAR fluorescence in NRVMs was greater upon faster electrical pacing compared to treatment with AngII, suggesting that CaMKII is more sensitive to direct changes in voltage than to hormonal signaling in these cells. Between muscle cell types, we found that CaMKAR in skeletal myocytes demonstrated a similar maximum increase in fluorescence compared to cardiomyocytes. Notably, skeletal myocytes were paced at a higher frequency than cardiomyocytes in our experiments.

Interestingly, we found in skeletal muscle that there appears to be an optimal rate of stimulation at which CaMKII is most efficiently activated, with 140 Hz stimulation showing a slower activation curve compared to 70 Hz. This phenomenon may be explained by a frequency-dependent mechanism wherein calcium release through the ryanodine receptor 1 (RyR1) becomes inactivated at rapid rates of stimulation [26, 27]. Furthermore, imaging CaMKAR on the timescale of milliseconds provides the granularity to delineate discrete activation spikes of neuronal CaMKII, suggesting the important role of CaMKII activation within individual action potentials.

Our efforts at modeling CaMKII kinetics in these systems also reveals CaMKAR’s ability to generate data that is sufficient to inform inferences about CaMKII regulation at the molecular level. Our computational framework for CaMKII-CaM-Ca^2+^ equilibriums reveals different rates of accumulation of active-CaMKII in each cell type in a manner that is also frequency-dependent. These CaMKII models that are based on real-time signal data from CaMKAR will enhance how we make predictions about the ways in which CaMKII systems might respond to cellular stressors and disequilibria that are not easily reproduced and tested at the wet bench. In this way, CaMKAR not only expands our ability to produce high-quality experimental data of CaMKII signaling in living cells but can also inform the design of future experiments before they have even been conducted.

Taken together, our signal measurements across cardiac and extracardiac tissue delineate, with previously unattainable precision, the range of temporal dynamics over which CaMKII operates to encode transient calcium signaling in different cell types. One conceivable mechanism by which these cell types achieve such variation in temporal dynamics lies in the differences across properties and subcellular localization of CaMKII isoforms and splice variants. Therefore, further investigation aimed at identifying the factors driving this observed variability will no doubt benefit from CaMKAR’s ability to retain its properties when targeted to the nucleus, cytoplasm, or mitochondria. Other important factors that could potentially drive CaMKII to unique kinetic phenotypes include post-translational modifications such as phosphorylation, oxidation, or O-GlcNAcylation, protein-protein interactions, and cell-dependent Ca^2+^ signaling.

Each of our models here involve measuring CaMKII kinetics in healthy cell systems. However, CaMKAR will also allow us to conduct future experiments aimed at identifying unique features of CaMKII activation patterns in disease models, including atrial fibrillation and heart failure, where CaMKII is known to drive pathology.

As CaMKII becomes increasingly implicated in the pathogenesis of various disease states, it is becoming more important than ever to develop a precise understanding of CaMKII’s unique enzymatic profile across its different physiologic and pathologic roles in humans. Considering what we have demonstrated, we posit that the CaMKAR sensor will play a crucial role in this endeavor. We believe that CaMKAR is a powerful tool with the ability to aid the search for CaMKII inhibitor candidates in cardiac tissue and also help validate potential candidates in order to minimize their extracardiac impacts.

## Methods

### Plasmids and molecular biology

PcDNA3.1-CMV-CaMKAR was modified with site directed mutagenesis (KDL mix, NEB) using oligonucleotides encoding modifications to either N or C terminus of CaMKAR in order to generate localized variants of CaMKAR. The jRCaMP1a calcium biosensor was used directly from pGP-CMV-NES-jRCaMP1a, gifted by Douglas Kim & GENIE Project (Addgene plasmid # 61562;http://n2t.net/addgene:61562; RRID:Addgene_61562) [28] (27). Human CaMKIIδ was used to construct CaMKII splice variants and then mutated to include or delete the exons represented in the several splice variants (KDL mix, NEB). Similarly, amino acids 1-69 of SU9 were added to the N-terminus of CaMKIIδC to create mitoCaMKII. CaMKAR, jRCAMP1a, and CaMKII variants were encoded into lentiviruses.

### Cell Culture

Neonatal rat ventricular myocytes (NRVMs) were isolated and cultured as previously described [29]. Briefly, day 1 to 2 newborn Sprague-Dawley rats (Envigo) were placed in an anesthesia chamber with a paper towel that had been saturated with isoflurane. Rats were then sterilized with ethanol. The heart was removed via midline incision and placed in Krebs-Henseleit buffer to be mechanistically dissociated. The disaggregated myocytes were then cultured in 48-and 24-well plates and incubated for 1 to 2 days prior to use. Isolated myocytes were cultured in DMEM (Gibco) supplemented with 10% FBS (Gibco) and Pen/Strep (Gibco).Maintenance: HEK293T/17 cells (ATCC CRL-11268) were maintained in DMEM (L-glutamine, Sodium Pyruvate, Non-essential amino acids; Gibco) supplemented with 10% FBS (Gibco) and Pen/Strep (Gibco) between 10%-95% confluence. Lentivirus production and infection: HEK293T/17 cells were seeded into 10cm dishes at 400k cells/mL. These cells were transfected using TransIT-Lenti (Mirus Bio) using a ratio of 5:3.75:1.25 of packaging plasmid:psPAX2:pMD2.G yielding a total of 10 µg per dish. After 48 hours, supernatant was collected, clarified, and concentrated 10-fold using Lenti-X concentrator (Takara). Lentivirus aliquots were stored at-80 °C until functional titration and use. Infection occurred in the presence of 10 μg/mL polybrene reagent (Sigma).

Plasmid Transfection: HEK293T/17 cells were plated into PDL-coated 24-well plates at 400k cells/mL. Each well was transfected with 500 ng of DNA complexed with 1 µL of JetPrime reagent according to Polyplus JetPrime protocol. Cells were examined 24-48 hrs post transfection. Adenovirus infection: CMV-CaMKAR-encoding Ad5 adenovirus was synthesized directly by Vector Biolabs. NRVMs were infected 24 hours post isolation at multiplicity of infection 20. Cells were analyzed 48 hours post infection.

### CaMKAR imaging in NRVMs

Timelapse microscopy was performed using an Olympus IX-83 inverted widefield microscope equipped with an ORCA Flash 4.0 and Lumencor SOLA light source. CaMKAR signal was captured at 200 ms exposure using the following channels: excitation filters ET402/15x and ET490/20x and emission filter ET525/35m (Chroma Technology). CaMKAR Signal (R) is defined as the ∼488-nm-excited intensity divided by the ∼400-nm-excited intensity. Calcium imaging was collected in the TRITC channel ex 555/em. 590 nm. Confocal imaging was performed using a Zeiss LSM880 Airyscan FAST. Using excitation lasers 405 nm and 488 nm and collecting emission window at 520 ± 10 nm. Image analysis: Otsu segmentation was used to track individual cells and their intensity in 488 nm and 400 nm channels at every timepoint in CellProfiler. Individual cell and well values were imported to R Studio for tabulation and summary statistics including mean, standard deviation, and n calculation.

### CaMKAR imaging in skeletal muscle

All the animal procedures and protocols were reviewed and approved by the Institutional Animal Care and Use Committees of the University of Maryland. Male C57BL/6J mice (Charles River, Wilmington, MA) were used. All mice used (9 mice) were between 30-60 days of age. Environmental conditions were maintained with a 12-h light/dark cycle and constant temperature (21–23°C) and humidity (55 ± 10%). The cages contained corncob bedding (Harlan Teklad 7902) and environmental enrichment (cotton nestlet). Mice were supplied with dry chow (irradiated rodent diet; Harlan Teklad 2981) and allowed water ad libitum. Electroporation was carried out on 4-week-old C57BL mice (2). Subcutaneous injection of 20-30 μL of hyaluronidase (2.5 mg/ml) was administered into the plantar pad of a mouse anesthetized with 3%-4.5% isoflurane in O₂ (1 L/min), using a 33-gauge needle. One hour later the mouse was again anesthetized and ∼60-70 μg of CAMKAR plasmid DNA were injected into the footpad. The animal was kept on an isothermal pad for 5 minutes; then, two surgical stainless-steel electrodes were placed subcutaneously close to the proximal and distal tendons of the flexor digitorum brevis (FDB) muscle and 20 pulses of 100 V/cm, 20 ms in duration, were applied at 1 Hz via a commercial high current capacity output stage (ECM 830, BTX, Harvard Apparatus, Holliston, MA).

One week later, single muscle fibers were enzymatically dissociated from the injected FDB muscles and cultured as described below [30]. FDB muscle was isolated from male adult mice, enzymatically dissociated with collagenase type I (Sigma-Aldrich, St. Louis, MO) in spinner minimum essential medium (S-MEM; Cat. No. 11380-037; Gibco, Carlsbad, CA) with 10% FBS (Cat. #100–106; Gemini Bio-Products, West Sacramento, CA), and 50 µg/ml gentamicin for 3-4 hours at 37° C. Muscle was then gently triturated to separate fibers in S-MEM with FBS 10% and gentamicin. Fibers were plated in MEM culture media with 2% FBS on glass-bottomed dishes (Matek Cor. Ashland, MA, Cat. No. P35G-1.0-14-C,) coated with laminin (Thermo Fisher, Rockford, IL, Cat. No. 23017-015). Fibers were maintained in culture for 2 to 20 h at 37°C, 5% CO2 before the experiments.

Positively transfected fibers were identified by the fluorescence expression profile excited with 488 nm and emitted light collected with a long-pass filter >510 nm. Cultured skeletal muscle FDB fibers expressing different CaMKARs constructs were imaged in 2 mL of L-15 media (ionic composition in mM: 137 NaCl, 5.7 KCl, 1.26 CaCl2, 1.8 MgCl2, pH 7.4; Life Technologies, Carlsbad, CA). All single fiber recordings were performed at room temperature, 22°C. Individual muscle fibers expressing CAMKAR were excited sequentially with a 488 nm laser, and the fluorescence emitted >505 mm was acquired a confocal system LSM 5 Live system (Carl Zeiss, Jena, Germany) mounted on a Zeiss Axiovert 200M inverted microscope. Imaging was performed in frame scan xy mode with images acquired at a rate of 4 frames per second (fps) for 105 s, using a 20x 0.5 N.A. dry or 63x 1.2 N.A. water immersion objective.

Electrical field stimuli were applied via two parallel platinum wires closely spaced (5 mm). Each stimulating pulse (1 ms, 10-20 V/cm) alternated the polarity using a custom pulse generator at 10, 70 or 140 Hz for 200 ms every 2 sec for 1 min. The average fluorescence intensity within the selected optical field in the distal segment of the fiber was recorded and background corrected by subtracting an average value recorded outside the cell. To estimate CAMKAR activity, the first 20 images of each fiber were used as a control to establish the resting steady-state fluorescence level (F0). The mean fluorescence intensity within the selected region of interest (ROI) in each myofiber was measured using ImageJ software (National Institutes of Health, Bethesda, MD). The mean F0 value for each ROI was used to normalize the activity signals as ΔF/F0. To avoid one source of systematic bias, experimental and control measurements were alternated whenever possible and concurrent controls were always performed. Data spreadsheets were generated from the raw fluorescent images. Data were initially processed in Excel (Microsoft, Redmond, WA, USA). Visual Basic (Microsoft, Redmond, WA, USA) macros were used to systematically determine the CAMKAR fluorescence ratio (488/405). Data were then analyzed and plotted using OriginPro 2022 (OriginLab Corporation, Northampton, MA, USA).

### CaMKAR imaging in neurons

Rat hippocampal neurons were cultured as previously described [31]. Briefly, hippocampal neurons obtained from embryonic day 18 Sprague-Dawley rats were plated onto poly-l-lysine-coated glass coverslips and cultured in neurobasal based media. Neurons were transfected with jRGECO1a and pCAG-CaMKAR plasmids using Lipofectamine 2000 (Invitrogen) at 15-18 days in vitro and imaged 2-4 days later. All experimental procedures involving animals were conducted according to the National Institutes of Health guidelines for animal research and were approved by the Animal Care and Use Committee at Johns Hopkins University School of Medicine. Imaging was performed on a Zeiss spinning-disk confocal microscope equipped with an AxioObserver Z1 stand, a Yokogawa CSU-X1A 5000 spinning-disk unit, Photometrics Evolve EMCCD camera, four lasers (405/488/561/639 nm) and a Definite Focus system. Excitation-ratiometric imaging of CaMKAR was performed using the 488-and 405-nm laser lines, a RQFT 405/488/568/647 dichroic mirror and a 525/50 emission filter. jRGECO1a was imaged using the 561-nm laser, the RQFT 405/488/568/647 dichroic mirror and a 629/62 emission filter.

The glycine-stimulation was performed as previously described [31]. Neuron culture media were supplemented with 200 μM DL-AP5 1-2 days before experiments. On the day of experiments, neurons were preincubated in basal artificial cerebrospinal fluid (ACSF, 120 mM NaCl, 5 mM KCl, 2 mM CaCl2, 1 mM MgCl2, 10 mM d-glucose and 10 mM HEPES, pH 7.4, supplemented with 200 μM DL-AP5, 1 μM TTX, 1 μM strychnine and 100 μM picrotoxin) for at least 1 h before imaging. Images (single Z slice) were acquired in streaming mode (∼1.2 Hz). Neurons were recorded for 5-10 minutes at baseline and then were stimulated with glycine (ACSF minus MgCl_2_, supplemented with 1 μM TTX, 1 μM strychnine, 100 μM picrotoxin and 200 μM glycine). Calcium transients were identified and analyzed as previously reported [31].

### Statistics and schematics

Imaging summary data condensed and organized with R Studio. Statistical testing done with GraphPad Prism v8.2.0 as described in each figure legend.

### CaMKII Kinetic Modeling

The dynamics of CaMKII, CaM, and calcium concentrations were described as a set of nonlinear ordinary differential equations (ODEs) derived from the law of mass action. CaMKII and calmodulin state transitions were described by biochemical reactions and rate equations. The CaMKII and CaM model was adapted and modified from the previously proposed CaMKIIα kinetics models [21, 32, 33]. The equations governing CaMKII dynamics represent its various states, including free and bound states, as well as phosphorylated and dephosphorylated states. These equations include the binding of CaM and calcium ions to CaMKII, and phosphorylation and dephosphorylation processes.

The equation for calcium dynamics was described by a waveform function. For the modeling of calcium binding to CaM, a previously established scheme [33] was employed and simplified. Four different Ca2+/CaM complexes (CaM0, CaM2N, CaM2C, and CaM4), four different Ca2+/CaM-CaMKII (K) complexes, and four different Ca2+/CaM-CaMKIIP (P) complex states were presented, assuming two or four calcium bind to N lobe or C lobe of CaM subunit.

The computational framework was implemented using the Julia programming language. The reaction network was defined and constructed using the Catalyst.jl and ModelingToolkit.jl packages in Julia. It includes reactions for the binding of calcium ions to CaM, the formation of CaM-CaMKII complexes, phosphorylation and dephosphorylation of CaMKII. The DifferentialEquations package in Julia was used to solve the system of ODEs numerically. The initial value problem was solved using a numerical solver with high precision settings (absolute tolerance = 1e-9, relative tolerance = 1e-9), and the solution was saved at intervals of 0.0005 (save at = 0.0005) for analysis.

## Acknowledgments

We wish to acknowledge Chin-Shun Wu for establishing the CaMKII framework and computing code employed in CaMKII kinetic modeling. We also wish to thank R. C. Johnson for helping with subcloning.

---

**Table S1.** Parameter Description.

| Parameter | Description | Skeletal Muscle | Cardiac NRVM | Neuron<sup>[20]</sup> |
| --- | --- | --- | --- | --- |
| CaMT (M) | Total CaM concentration | $3 \times 10^{-5}$ | $3 \times 10^{-5}$ | $3 \times 10^{-5}$ |
| CaMKII_T (M) | Total CaMKII concentration | $7 \times 10^{-5}$ | $7 \times 10^{-5}$ | $7 \times 10^{-5}$ |
| decay_CaM (s) | CaM on/off from CaMKII-P (PCaMi_off) | 3 | 3 | 3 |
| k<sub>1C_on</sub> (μM<sup>-1</sup>s<sup>-1</sup>) | Ca<sup>2+</sup> binding to CaM (first C-lobe) | $5 \times 10^6$ | $3 \times 10^6$ | $5 \times 10^6$ |
| phospho_rate (s<sup>-1</sup>) | k<sub>phos_CaMK</sub> (CaM-CaMKII → CaM-CaMKIIP) | 1 | 0.2 | 1 |
| phosphatase (s<sup>-1</sup>) | k<sub>dephospho</sub> (CaMKIIP → CaMKII) | 0.3 | 0.3 | 1 |

## References

1. Frankland, P.W., et al., Alpha-CaMKII-dependent plasticity in the cortex is required for permanent memory. Nature, 2001. 411(6835): p. 309-13.

2. Opazo, P., et al., CaMKII triggers the diffusional trapping of surface AMPARs through phosphorylation of stargazin. Neuron, 2010. 67(2): p. 239–52.

3. Tomita, S., et al., Bidirectional synaptic plasticity regulated by phosphorylation of stargazin-like TARPs. Neuron, 2005. 45(2): p. 269–77.

4. Gaido, O.E.R., et al., CaMKII as a Therapeutic Target in Cardiovascular Disease. Annu Rev Pharmacol Toxicol, 2022.

5. Wu, Y., et al., Calmodulin kinase II is required for fight or flight sinoatrial node physiology. Proc Natl Acad Sci U S A, 2009. 106(14): p. 5972–7.

6. Luczak, E.D., et al., Mitochondrial CaMKII causes adverse metabolic reprogramming and dilated cardiomyopathy. Nat Commun, 2020. 11(1): p. 4416.

7. Wang, Q., et al., CaMKII oxidation is a critical performance/disease trade-off acquired at the dawn of vertebrate evolution. Nature Communications, 2021. 12(1): p. 3175.

8. Witczak, C.A., et al., CaMKII regulates contraction-but not insulin-induced glucose uptake in mouse skeletal muscle. Am J Physiol Endocrinol Metab, 2010. 298(6): p. E1150–60.

9. Lou, L.L. and H. Schulman, Distinct autophosphorylation sites sequentially produce autonomy and inhibition of the multifunctional Ca2+/calmodulin-dependent protein kinase. J Neurosci, 1989. 9(6): p. 2020–32.

10. Meyer, T., et al., Calmodulin trapping by calcium-calmodulin-dependent protein kinase. Science, 1992. 256(5060): p. 1199-202.

11. Gaido, O.E.R., et al., Novel Biosensor Identifies Ruxolitinib as a Potent and Cardioprotective CaMKII Inhibitor. bioRxiv, 2022: p. 2022.09.24.509320.

12. Sepulveda, M., et al., Role of CaMKII and ROS in rapid pacing-induced apoptosis. J Mol Cell Cardiol, 2013. 63: p. 135–45.

13. Wood, B.M., et al., Cardiac CaMKII activation promotes rapid translocation to its extra-dyadic targets. J Mol Cell Cardiol, 2018. 125: p. 18–28.

14. De Koninck, P. and H. Schulman, Sensitivity of CaM kinase II to the frequency of Ca2+ oscillations. Science, 1998. 279(5348): p. 227-30.

15. Li, H., et al., Calmodulin kinase II is required for angiotensin II-mediated vascular smooth muscle hypertrophy. Am J Physiol Heart Circ Physiol, 2010. 298(2): p. H688–98.

16. Palomeque, J., et al., Angiotensin II-Induced Oxidative Stress Resets the Ca2+ Dependence of Ca2+-Calmodulin Protein Kinase II and Promotes a Death Pathway Conserved Across Different Species. Circ Res, 2009. 105(12): p. 1204–1212.

17. Wagner, S., et al., NADPH oxidase 2 mediates angiotensin II-dependent cellular arrhythmias via PKA and CaMKII. J Mol Cell Cardiol, 2014. 75: p. 206–15.

18. Robison, P., E.O. Hernandez-Ochoa, and M.F. Schneider, Atypical behavior of NFATc1 in cultured intercostal myofibers. Skelet Muscle, 2014. 4(1): p. 1.

19. Urakubo, H., et al., In vitro reconstitution of a CaMKII memory switch by an NMDA receptor-derived peptide. Biophys J, 2014. 106(6): p. 1414–20.

20. Michalski, P.J., First demonstration of bistability in CaMKII, a memory-related kinase. Biophys J, 2014. 106(6): p. 1233–5.

21. Chang, J.Y., et al., Mechanisms of Ca(2+)/calmodulin-dependent kinase II activation in single dendritic spines. Nat Commun, 2019. 10(1): p. 2784.

22. Duran, J., et al., CaMKIIdelta Splice Variants in the Healthy and Diseased Heart. Front Cell Dev Biol, 2021. 9: p. 644630.

23. Gaertner, T.R., et al., Comparative analyses of the three-dimensional structures and enzymatic properties of alpha, beta, gamma and delta isoforms of Ca2+-calmodulin-dependent protein kinase II. J Biol Chem, 2004. 279(13): p. 12484–94.

24. Zalcman, G., N. Federman, and A. Romano, CaMKII Isoforms in Learning and Memory: Localization and Function. Front Mol Neurosci, 2018. 11: p. 445.

25. Tobimatsu, T. and H. Fujisawa, Tissue-specific expression of four types of rat calmodulin-dependent protein kinase II mRNAs. J Biol Chem, 1989. 264(30): p. 17907–12.

26. Laver, D.R. and G.D. Lamb, Inactivation of Ca2+ release channels (ryanodine receptors RyR1 and RyR2) with rapid steps in [Ca2+] and voltage. Biophys J, 1998. 74(5): p. 2352–64.

27. Hernandez-Ochoa, E.O., et al., Critical Role of Intracellular RyR1 Calcium Release Channels in Skeletal Muscle Function and Disease. Front Physiol, 2015. 6: p. 420.

28. Dana, H., et al., Sensitive red protein calcium indicators for imaging neural activity. Elife, 2016. 5.

29. Mishra, S., et al., Inhibition of phosphodiesterase type 9 reduces obesity and cardiometabolic syndrome in mice. J Clin Invest, 2021. 131(21).

30. Bibollet, H., et al., Functional Site-Directed Fluorometry in Native Cells to Study Skeletal Muscle Excitability. J Vis Exp, 2023(196).

31. Zhang, J.F., et al., An ultrasensitive biosensor for high-resolution kinase activity imaging in awake mice. Nat Chem Biol, 2021. 17(1): p. 39–46.

32. Lucic, V., G.J. Greif, and M.B. Kennedy, Detailed state model of CaMKII activation and autophosphorylation. Eur Biophys J, 2008. 38(1): p. 83–98.

33. Pepke, S., et al., A dynamic model of interactions of Ca2+, calmodulin, and catalytic subunits of Ca2+/calmodulin-dependent protein kinase II. PLoS Comput Biol, 2010. 6(2): p. e1000675.

